# Phenotypically distinct human sequence is widespread in publicly archived microbial reads: an evaluation of methods for its detection

**DOI:** 10.1101/857508

**Authors:** Stephen J. Bush, Thomas R. Connor, Tim E. A. Peto, Derrick W. Crook, A. Sarah Walker

## Abstract

Sequencing data from host-associated microbes can often be contaminated by the body of the investigator or research subject. Human DNA is typically removed from microbial reads either by subtractive alignment (dropping all reads that map to the human genome) or using a read classification tool to predict those of human origin, and then discarding them. To inform best practice guidelines, we benchmarked 8 alignment-based and 2 classification-based methods of human read detection using simulated data from 10 clinically prevalent bacteria and 3 viruses, into which contaminating human reads had been added.

While the majority of methods successfully detected > 99% of the human reads, they were distinguishable by variance. The most precise methods, with negligible variance, were Bowtie2 and SNAP, both of which misclassified few, if any, bacterial reads (and no viral reads) as human. While correctly detecting a similar number of human reads, methods based on taxonomic classification, such as Kraken2 and Centrifuge, often misclassified bacterial reads as human, the extent of which was species-specific. Among the most sensitive methods of human read detection was BWA, although this also made the greatest number of false positive classifications. Across all methods, the set of human reads not identified as such, although often representing < 0.1% of the total reads, were non-randomly distributed along the human genome with many originating from the repeat-rich sex chromosomes.

For viral reads and longer (> 300bp) bacterial reads, the highest performing approaches were classification-based, using Kraken2 or Centrifuge. For shorter (150-300bp) bacterial reads, combining multiple methods of human read detection maximised the recovery of human reads from contaminated short read datasets without being compromised by false positives. The highest-performing approach with shorter bacterial reads was a two-stage classification using Bowtie2 followed by SNAP. Using this approach, we re-examined 11,577 publicly archived bacterial readsets for hitherto undetected human contamination. We were able to extract a sufficient number of reads to call known human SNPs, including those with clinical significance, in 6% of the samples. These results show that phenotypically-distinct human sequence is widespread in publicly-archived (and nominally pure) bacterial datasets.

## Background

Sequencing data from host-associated microbes, including metagenomic read sets, can often be contaminated by the body of the investigator or research subject (1). Furthermore, as the human genome is around 1000-fold larger than most bacterial genomes, sequencing a tissue biopsy containing equal numbers of human and bacterial cells would still produce a sample containing 99.9% human DNA (2). Accordingly, (human) contaminants need to be removed prior to downstream analysis (3) so as to minimise false-positive associations and reduce batch effects. The incomplete removal of human reads from nominally pure microbial datasets could theoretically lead to large volumes of residual human DNA being deposited in public archives. This raises numerous ethical concerns (4), particularly as individuals can be distinguished even within large, pooled, genomic datasets (5). This is not just a theoretical problem: previous studies have identified substantial cross-species contamination in genome assemblies (6), including human DNA comprising 2% of the purportedly complete *Plasmodium gaboni* assembly, a known human parasite (7), and 492 of 2749 non-primate assemblies containing the primate-specific AluY element (8).

With ever-increasing volumes of genomic (and metagenomic) data being deposited in public archives, there is a practical need to benchmark methods of human read detection so as to inform best practice guidelines. These methods follow two basic approaches: subtractive alignment and direct classification. For the former, reads are mapped to the human genome using a short read aligner such as BWA-mem (as in (9, 10)), Bowtie2 (as in (11, 12)) or SNAP (as in (13), or as part of the SURPI – Sequence-based Ultra-Rapid Pathogen Identification – pipeline (14)) with successfully mapped reads then subtracted from the dataset. Numerous pre-processing tools have been developed using this approach including CS-SCORE (15), DeconSeq (16), GenCoF (17), and MetaGeniE (18) (which employ BWA-fastmap, BWA-sw, Bowtie2, and both BWA-mem and Bowtie2, respectively). The second approach is to classify, and then discard, human content by predicting the taxonomic origin of each read using a database of human, bacterial and viral genomes, such as RefSeq or NCBI nr, using k-mer based classification tools such as Centrifuge (19) or Kraken (20), as in two studies employing the latter (2, 21).

However, no comparisons to date have identified the optimal method. We therefore evaluated several variations on these two basic approaches: by mapping all reads within a mixed dataset to the human genome, and by predicting read origin with the taxonomic classifiers Centrifuge and Kraken, using both all-species and human-only databases. When mapping reads to the human genome, we used eight commonly-employed aligners – Bowtie2 (22), BWA-mem (23), GEM (24), HISAT2 (25), minimap2 (26), Novoalign (www.novocraft.com), SMALT (http://www.sanger.ac.uk/science/tools/smalt-0), and SNAP (27) – alongside two different versions of the human primary assembly, GRCh38 and GRCh37, the latter of lower quality. We included the lower quality (and hence less accurate/complete) assembly because in real data we expect contaminating sequences could be degraded and would not necessarily precisely map (for example, gaps in the GRCh37 assembly are apparent around the centromeres and short arms of acrocentric chromosomes (28)). In total, this represents 16 different approaches to subtractive alignment (8 aligners * 2 genomes) and 4 different approaches to direct classification (2 classifiers * 2 databases), 20 methods in total.

To evaluate each method, we simulated 3 sets of 150bp and 3 sets of 300bp paired-end reads (characteristic of the Illumina NextSeq and MiSeq platforms, respectively) at 10-fold coverage from 10 clinically common bacteria with fully sequenced (closed) core genomes: the Gram-positive *Clostridioides difficile*, *Listeria monocytogenes, Staphylococcus aureus*, and *Streptococcus pneumoniae,* the Gram-negative *Escherichia coli*, *Klebsiella pneumoniae, Neisseria gonorrhoeae, Salmonella enterica,* and *Shigella dysenteriae*, and *Mycobacterium tuberculosis*. These genomes were used in a previous benchmarking study assessing the performance of bacterial SNP calling pipelines (29). To each of these bacterial readsets we added human reads equivalent to between 1 and 10% of the bacterial reads, at increments of 1%. We supplemented this data with 3 sets of 150bp and 3 sets of 300bp paired-end reads simulated at 100-fold coverage from 3 viral genomes (hepatitis C, HIV, influenza A), with human reads also added at 1-10% increments.

As we aimed to establish the optimal method for maximising the removal of human reads, we report, for each method, the number of true positive (i.e. human reads identified as human), false positive (microbial reads identified as human) and false negative (human reads not identified as such) classifications, alongside the summary metrics precision (positive predictive value, the proportion of reads classified as human that truly are human), recall (sensitivity) and F-score, the harmonic mean of precision and recall (30). We also characterise those regions of the human genome more likely to be unidentified and which would be inadvertently (and disproportionately) retained in an otherwise ‘human depleted’ microbial dataset. We were specifically interested in methods that minimised the number of false negative calls and so also evaluated two-stage approaches, testing the sequential use of different aligners or classifiers.

## Results and Discussion

### Comparing methods for detecting human reads in a contaminated microbial dataset

The performance of 12 common methods of human read detection (8 aligners + 2 read classifiers each with 2 databases) was first evaluated using reads simulated from 10 closed bacterial genomes and 3 viral genomes (**Supplementary Table 1**), to which were added reads simulated from human genome GRCh38. This comparison contained 9360 records, comprising 2 read lengths (150bp and 300bp) * 3 replicates * 13 species * 10 incremental additions of human reads (comprising 1-10% of the total number of bacterial or viral reads) * 12 methods (that is, 780 records per method, of which 600 were bacterial). The number of reads simulated and the performance statistics for each method – the percentage of reads correctly classified as human (‘true positive rate’), the percentage of human reads not classified as human (‘false negative rate’), and the F-score (harmonic mean of precision and recall) – are given in **Supplementary Table 2**, with distributions illustrated in **Figure 1A**.

**Figure 1.**
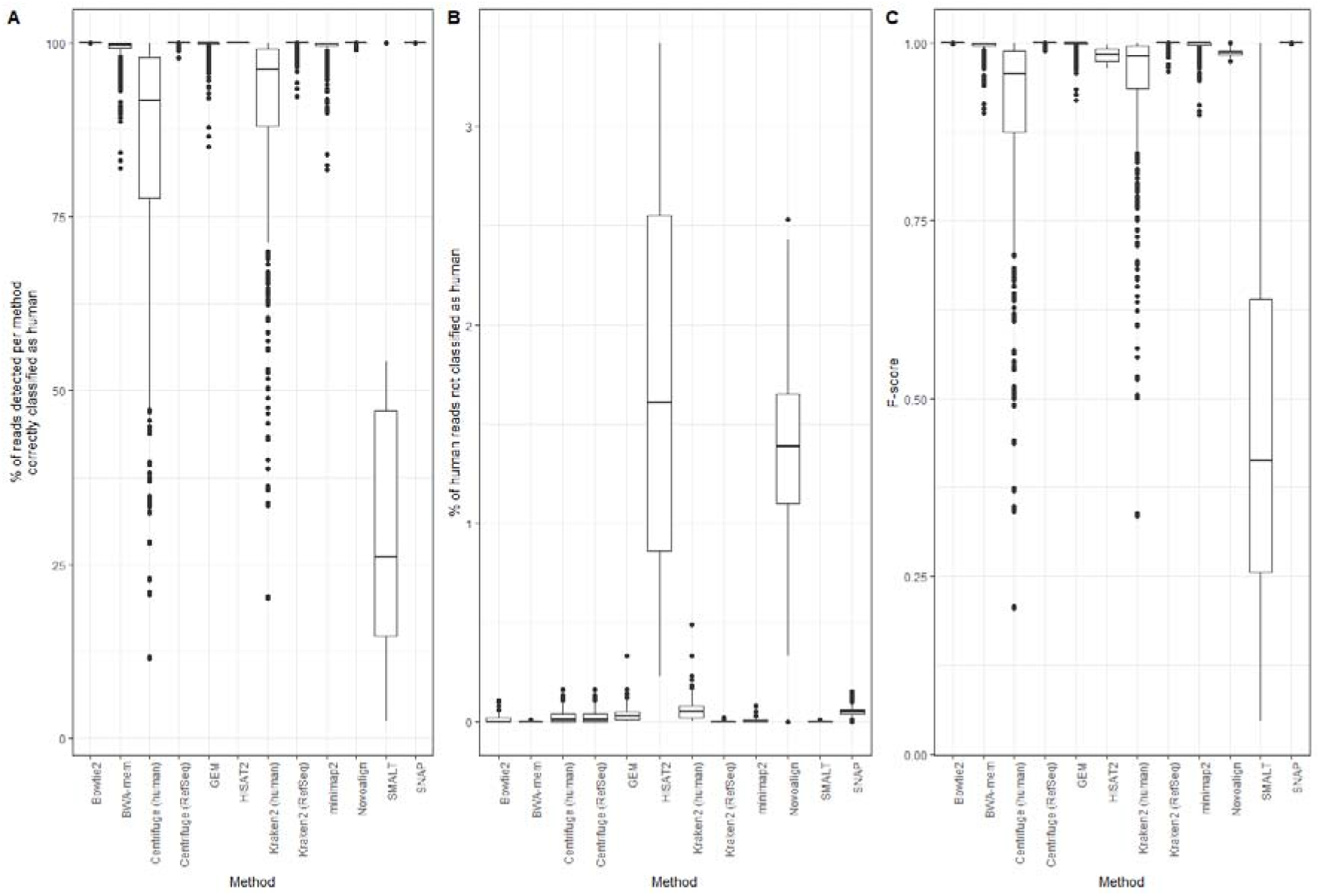
Performance of 12 different methods of identifying human reads within a range of microbial datasets (comprising 10 bacterial and 3 viral species sequenced at an average base-level coverage of 10- and 100-fold, respectively, each with between 1 and 10% simulated human contamination, using both 150bp and 300bp reads). All reads were simulated from, and where relevant aligned to, human genome version GRCh38.p12. The subfigures showc (A) the percentage of reads per method correctly classified as human, (B) the percentage of human reads not classified as human, and (C) the F-score. Data for this figure is available in **Supplementary Table 2**.

Although the median percentage of true positive classifications was high for the majority of methods, they were distinguishable by variance. This was only the case for bacteria as all methods using viral genomes identified 100% of the human reads (**Supplementary Table 2**). With bacteria, the highest true positive rates (> 99.9%), with negligible variance, were found using Bowtie2, HISAT2 or SNAP to align all reads against the human genome. The true positive rate was 100% in 582 of the 600 (bacterial) simulations using SNAP, 594 of the 600 simulations using Bowtie2, and all 600 simulations using HISAT2 (**Supplementary Table 2**). Similarly high true positive rates (around 99%), although with increasingly greater variance, were found when predicting human reads using a taxonomic classifier (Centrifuge/Kraken) with the RefSeq database (that is, the human genome plus prokaryotic genomes) and when using Novoalign, BWA-mem, GEM or Minimap2 to align all reads against the human genome. True positive rate was notably reduced (to < 90%), and variation substantially increased, when using either taxonomic classifier with a human-only database (**Figure 1A**). The poorest performing method (true positive rate < 25%) was the aligner SMALT which, although correctly identifying all human reads (that is, having a false negative rate of 0), could not reliably discriminate bacterial from human sequence and so made a large number of false positive calls (**Supplementary Table 2**).

Similarly, taxonomic classifiers using a complete genome database could discriminate between human and bacterial reads, whereas those using a human-only database made a far greater number of false positive calls (**Figure 1B**). This is because the human genome, as with other chordates, contains numerous genes of bacterial origin arising through horizontal gene transfer (HGT) (31), once thought of limited effect in animals due to their segregated germlines sheltering the genome from heritably meaningful exposure to foreign DNA (32). Accordingly, the proportion of false positive calls – while generally low – varied greatly by species (**Figure 2**). This was most notable for *C. difficile* (the genome of which is approx. 11% mobile genetic elements (33)) and *N. gonorrhoeae* (in which HGT from a human host has previously been characterised (34)) where the aligners BWA-mem and minimap2, and to a lesser extent GEM, made false positive calls at a rate > 5% in some simulations. Importantly, these methods misclassified bacterial reads as human even when no human reads were present in the dataset (**Supplementary Table 2**).

**Figure 2.**
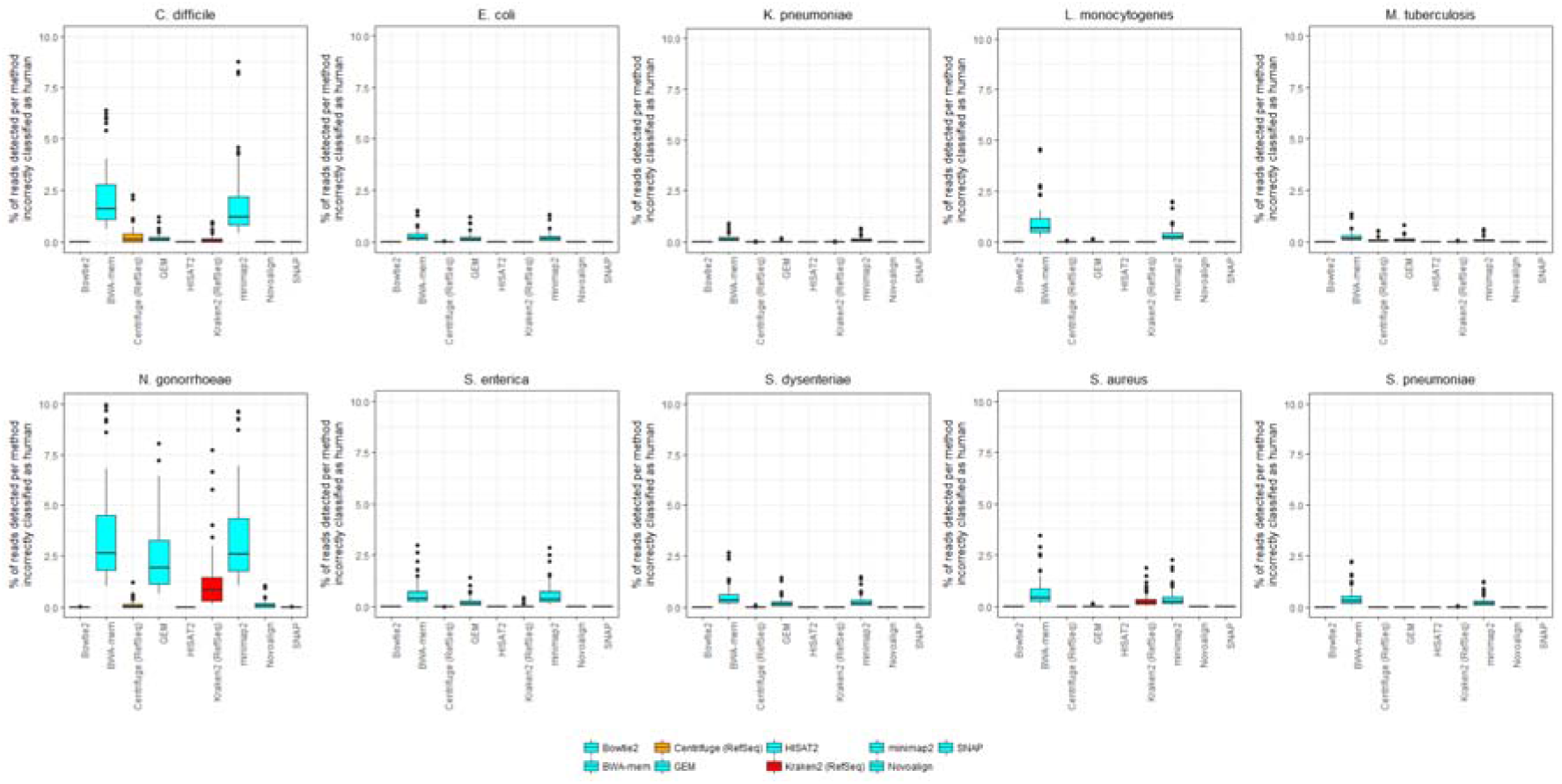
The percentage of reads incorrectly classified as human by 9 different methods of human read detection within a range of bacterial datasets, partitioned by species. Data for this figure is available in **Supplementary Table 2** and constitutes simulated reads at 10-fold coverage from each of 10 species supplemented with 0 to 10% human contamination, using both 150bp and 300bp reads. Data from 3 methods (the aligner SMALT and the classifiers Kraken2 and Centrifuge, each using a human-only database) are not shown. This is because these methods have a very high false positive rate across all species (**Figure 1**). Data from viral datasets is not shown because in our simulations no viral read was incorrectly classified as human, by any method (**Supplementary Table 3**).

Most methods produced a low number of false negative calls, of approximately < 1% of the total reads, with minimal variance (**Figure 1C**). This was particularly apparent for BWA-mem and Kraken2 (using the RefSeq database), for which the false negative rate using bacterial data was 0% for all 600 simulations, and for 573 of 600 simulations, respectively, with similar results seen with viral data (**Supplementary Table 2**). However, two aligners, HISAT2 and Novoalign, were prominent outliers, having relatively high false negative rates (with bacterial data, median 1.6% and 1.4%, respectively) and notably greater variance.

These results suggest that compared to the more-variable performance of a taxonomic classifier across the three summary metrics (Kraken and Centrifuge both make a number of false positive calls), the subtractive alignment of human reads is a generally higher-performing strategy for shorter (150-300bp) reads. Nevertheless, performance depends on choice of aligner. For example, HISAT2 had the highest true positive and lowest false positive rate (100% and 0%, respectively), but also made the greatest number of false negative calls (**Figure 1**). A lower number of false negative calls was made by, for example, BWA-mem, although at the expense of a higher false positive call rate. By contrast, Bowtie2 and SNAP had true positive and false positive rates comparable to HISAT2 (that is, approximating 100%), although far lower false negative rates.

Unlike the other aligners, the variance in false negative rate for Bowtie2 was related to read length. For 291 of the 300 bacterial simulations using 150bp reads, it was notable that false negative rate was 0% (**Supplementary Table 2**). However, and in clear contrast, for 288 of the 300 simulations using 300bp reads, false negative rate was > 0% (median 0.02%). The difference in call rate was because in this study, we made no attempt to optimise each aligner and so used only default parameters. Bowtie2 employs a ‘seed and extend’ alignment strategy, whereby exact matches of read substrings are used to ‘seed’ alignments of the full read, which by default is aligned in its entirety end-to-end. Read alignments are scored according to successful base matches and penalised for mismatches, with those meeting a minimum scoring threshold reported. It is important to note that Bowtie2 was originally designed to align shorter reads (< 300bp) with high sensitivity, with the scoring function optimised to that end. Without modifying the default parameters, longer reads would be slightly less likely to align, consistent with our observations.

To test the effect of read length on human read classification, we repeated the above simulations varying read length from 50bp to 1000bp, at 50bp increments (using the same parameters and formula for insert size as described in Materials and Methods). We repeated the simulations for the set of 10 bacterial genomes, although restricted analysis to 10 methods (we excluded the use of Centrifuge and Kraken with human-only databases) and to simulating a number of human reads equivalent to 10% of the number of bacterial reads. In total, this represents 6000 simulations (20 read lengths * 3 replicates * 10 species * 10 methods). Performance statistics are given in **Supplementary Table 3**, and the variance of F-score with read length illustrated in **Figure 3**. We excluded viral genomes from this analysis as the previous simulations showed that viral reads could be unambiguously distinguished from human reads. Accordingly, varying read length was not expected to have significant impact.

**Figure 3.**
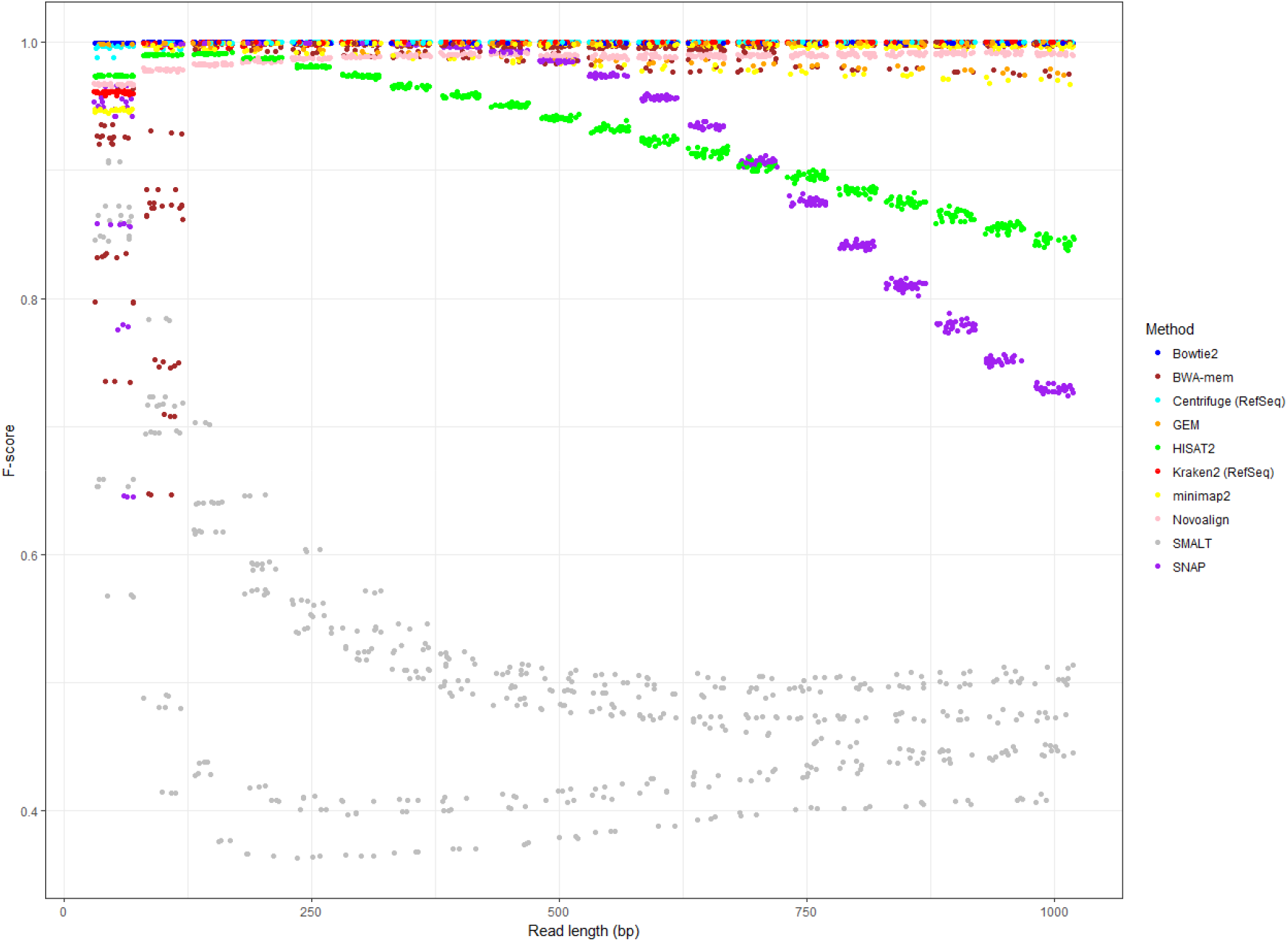
Performance of 10 different methods of identifying human reads within a range of bacterial datasets (comprising 10 bacterial species sequenced at an average base-level coverage of 10-fold, each with 10% simulated human contamination). All reads were simulated from, and where relevant aligned to, human genome version GRCh38.p12. Each point represents a simulation replicate, coloured according to method. Points are jittered to allow over-plotting. There is considerable overlap between points as many methods perform equivalently highly when using long reads. Data for this figure is available in **Supplementary Table 3**.

For longer (> 300bp) bacterial reads, the classification-based methods (Kraken and Centrifuge) consistently produced precision, recall and F-scores of 1 (**Supplementary Table 3**). By contrast, alignment-based methods were especially sensitive to read length since implicitly this affects the appropriateness of the parameters used to score alignments (the default parameters of most aligners assume reads of shorter length). All methods performed with comparable F-scores if using reads within the range of 150-300bp (i.e. read lengths expected of Illumina HiSeq/NextSeq/MiSeq sequencers), although there was considerable variance in performance at lower read lengths, with the exception of Bowtie2 (which performs highly even with 50bp reads). At higher read lengths, Bowtie2, HISAT2 and SNAP (when using default parameters) all declined in performance, with the F-scores of Bowtie2 and HISAT2 gradually declining from approx. 200bp and 500bp, respectively. SNAP declined in performance sharply when using reads greater than 400bp, although doing so is contrary to recommended use (to enable SNAP to process reads > 400bp, the value of MAX_READ_LENGTH needed to be manually edited in SNAPLib/Read.h before compiling). SMALT, if used with default parameters, appeared optimised for shorter (< 250bp) reads, and showed consistently poorer performance (plateauing at an approx. F-score of 0.5) for longer reads. SMALT was also strongly affected by species (see the highly variable F-score in **Figure 1C**), with multiple F-score distributions apparent in **Figure 3**.

We found no discernible difference in true or false positive classification rates when aligning reads to lower-quality human genomes (that is, older assemblies) (**Supplementary Figure 1**).

### All methods of human read depletion fail to classify reads from similar regions

We were particularly interested in false negative calls – human reads that were not classified as such and so would have been retained within a microbial dataset. In contrast, false positive calls are likely to be reads from human genes with microbial homology (and vice versa), the exclusion of which would artificially reduce the read depth of a microbial dataset, albeit only to a small degree.

By reference to the GRCh38.p12 and GRCh37.p13 gene coordinates (from Ensembl BioMart (35)), we identified, per method, the proportion of reads with false negative classifications (i.e., human reads not classified as human; hereafter ‘false negative reads’) deriving from different genomic regions (**Supplementary Table 4**). With Kraken and Centrifuge (when provided a combined microbial and human database), it was particularly notable that in absolute terms there were not only few false negative calls but few in genic regions (**Supplementary Table 4**). This is likely because both methods avoid (intrinsically inexact) alignments to make exact-match queries against a database of *k*-mers, with larger databases – and the uniqueness of gene sequences – affording greater resolving power. While Kraken classifies reads on the basis of the lowest common ancestor for all genomes containing its constituent *k*-mers (20), Centrifuge can also assign a single sequence to multiple taxonomic categories. However, as the present task is essentially one of binary classification – discriminating human from non-human reads – we found only a modest difference in performance between the two approaches. Using the RefSeq database, Kraken had greater variance in false positive calls than Centrifuge although compensated with fewer false negative calls (**Figure 1**). For both methods the presence of at least one or more unique *k*-mers among the reads sequenced from human genes appears sufficient to discriminate them from essentially all bacterial genomes (unique *k*-mers ostensibly reflecting divergent evolutionary histories).

**Table 1** summarises, per method, the number of unique genes from which one or more false negative reads were derived (hereafter, ‘false negative genes’), or to which one or more bacterial reads had been classified (hereafter, ‘false positive genes’), across all 13 species and all replicates of each method. Consistent with **Figure 1C**, virtually all human reads were detected by methods with zero or negligible false negative rate such as BWA-mem and SMALT, with reads undetected by Kraken2 or Centrifuge (when using the RefSeq database) representing < 40 genes. The number of genes represented by the two methods with the highest false negative (HISAT2) and false positive (SMALT) rates accounted for the majority of genes in the human genome (> 20,000), suggesting that reads misclassified by these methods were unlikely to have common characteristics. Accordingly, we would not expect these sets of genes to be significantly enriched for Gene Ontology (GO) terms if false negative, or false positive, calls were randomly distributed across the genome.

**Table 1.** Total number of genes, per method, whose reads could not be classified as human (that is, false negative calls) or to which microbial reads had been classified (that is, false positive calls). The number of unique genes per method is given across all species and all replicates of that method. Note that as the current Centrifuge and Kraken RefSeq-based databases contain human assembly GRCh38, it is unnecessary to create an equivalent with GRCh37.

| Method | No. of human genes whose reads could not be classified as human (i.e. false negative classifications) |  | No. of human genes to which bacterial reads had been classified (i.e. false positive classifications) |  |
| --- | --- | --- | --- | --- |
|  | GRCh38.p12 | GRCh37.p13 | GRCh38.p12 | GRCh37.p13 |
| Bowtie2 | 627 | 593 | 0 | 0 |
| BWA-mem | 0 | 0 | 13,170 | 587 |
| Centrifuge (human) | 23 | n/a | 0 | n/a |
| Centrifuge (RefSeq) | 25 | n/a | 0 | n/a |
| GEM | 42 | 64 | 120 | 110 |
| HISAT2 | 22,012 | 21,053 | 0 | 0 |
| Kraken2 (human) | 1,547 | n/a | 0 | n/a |
| Kraken2 (RefSeq) | 37 | n/a | 0 | n/a |
| Novoalign | 3,588 | 3,367 | 11 | 10 |
| minimap2 | 660 | 516 | 443 | 421 |
| SMALT | 0 | 2 | 35,518 | 33,874 |
| SNAP | 3,283 | 3,047 | 0 | 0 |

Nevertheless, we found that for the remaining methods (which were more sensitive and/or more precise) each set of false negative genes was enriched for a broadly consistent set of GO terms related to the nervous system (**Supplementary Table 5**). For example, when aligning reads to the GRCh38.p12 assembly, undetected reads were enriched for genes with process terms such as ‘dendrite development’ (among the set of genes with reads undetected by GEM) and ‘glutamatergic synaptic transmission’ (minimap2) and component terms such as ‘synaptic membrane’ (SNAP, GEM and minimap2), ‘postsynaptic membrane’ (Bowtie2), ‘neuromuscular junction’ (GEM), ‘neuron part’ (minimap2), ‘photoreceptor disc membrane’ (Kraken2) and ‘astrocyte projection’ (Centrifuge). Similar results were found when aligning reads to the GRCh37.p13 assembly, with undetected reads enriched for genes with process terms including ‘neural tube formation’ (Bowtie2) and ‘synapse assembly’ (both minimap2 and SNAP) (**Supplementary Table 5**). An equivalent set of nervous system-related GO terms was found for each set of false positive genes (that is, genes with a likely microbial homologue) in both assemblies. In this case, misclassified microbial reads were also enriched for genes with component terms such as ‘neuronal cell body’ (GEM; GRCh38.p12) and process terms such as ‘postsynaptic membrane assembly’ (BWA-mem; GRCh37.p13) (**Supplementary Table 6**).

These results suggest that in general, alignment-based methods of human read detection consistently fail to classify reads from functionally-similar genes. Further to this, a relatively high proportion of false negative reads originated from the sex chromosomes (**Figure 4** and **Supplementary Table 4**), both known for their high (>50%) repeat content (36, 37). This was particularly apparent when using Novoalign (from which the median percentage of false negative reads from chromosomes X and Y were 6.4% and 18.3%, respectively), and to a lesser extent Bowtie2 (6.0% X, 4.9% Y) and Centrifuge, when using the RefSeq database (13.8% X, 0% Y).

**Figure 4.**
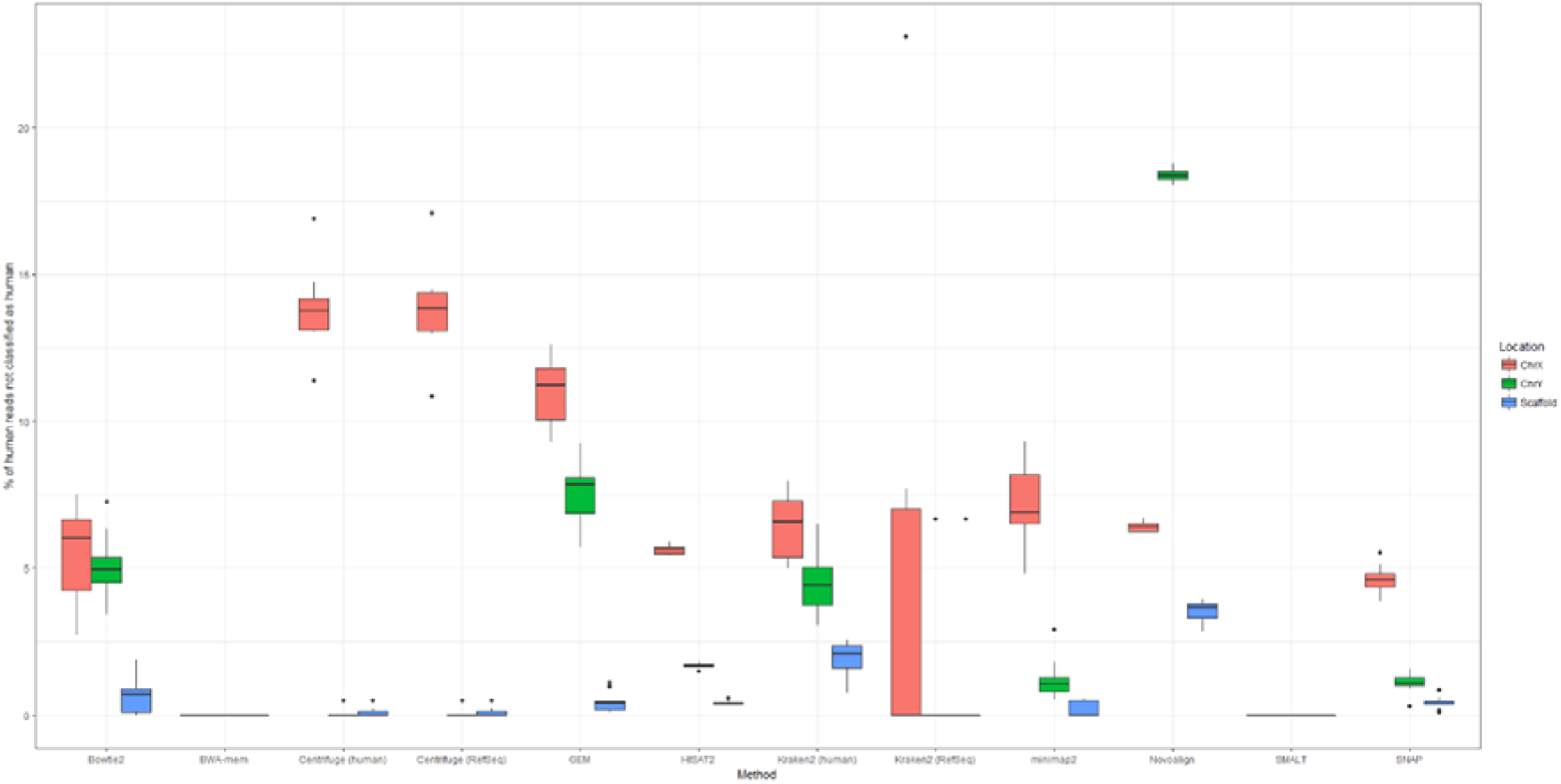
Proportion of human reads not classified as human by 9 different methods of human read detection, and their genomic location. Data for this figure is available in **Supplementary Table 4**.

It is important to note that it is impossible to align shorter sequencing reads across lengthy repeats. To resolve this problem, a previous study has advocated a Y-specific approach to read classification. RecoverY (38) is a tool for selecting Y-specific reads by reference to a database of *k*-mers built from single-copy genes on the Y chromosome. The original purpose of this was, assuming only human reads as input, to detect and remove non-Y reads which could otherwise confound a genome assembler. Whilst in theory this approach could be combined with others tested here, in practice several of the methods did not suffer from sex chromosome specific mapping issues and would hence be preferable.

Overall, the non-random distribution of false negative reads suggests that many methods of human read detection failed to detect reads from similar regions. It follows that a possible strategy for maximising the number of correctly identified human reads is to combine approaches so that multiple methods could compensate for locally-poor mapping.

### Human (short) read detection is improved by combining results from multiple aligners or classifiers

The retention of false negative reads – human reads not identified as such – within a nominally ‘pure’ microbial dataset is an ethical and practical concern. We reasoned that for contaminated short (150-300bp) bacterial readsets, for which human reads are not consistently removed (**Supplementary Table 2**), a two-stage approach – the sequential use of different aligners/classifiers – could map a greater number of reads to the human genome without losing specificity. Accordingly, we re-calculated true positive, false positive and false negative rate after combining the set of classifications made by any two methods. This two-stage approach first classified, and then discarded, ‘human’ reads using one method, and then performed a second round of classification using a second method. In this way, a method with high precision could be supplemented by a method with high sensitivity, maximising the utility of both. For this analysis we excluded three approaches with high positive rates (**Figure 1**) that were unlikely to provide much added-value (SMALT, and human-only databases with Centrifuge and Kraken). We also restricted this analysis only to bacterial data because our simulations showed that individual methods could discriminate viral from human reads (**Supplementary Table 2**). The number of reads classified by each pairwise combination of methods are given in **Supplementary Table 7**, with F-score and false negative rate distributions illustrated in **Figure 5**.

**Figure 5.**
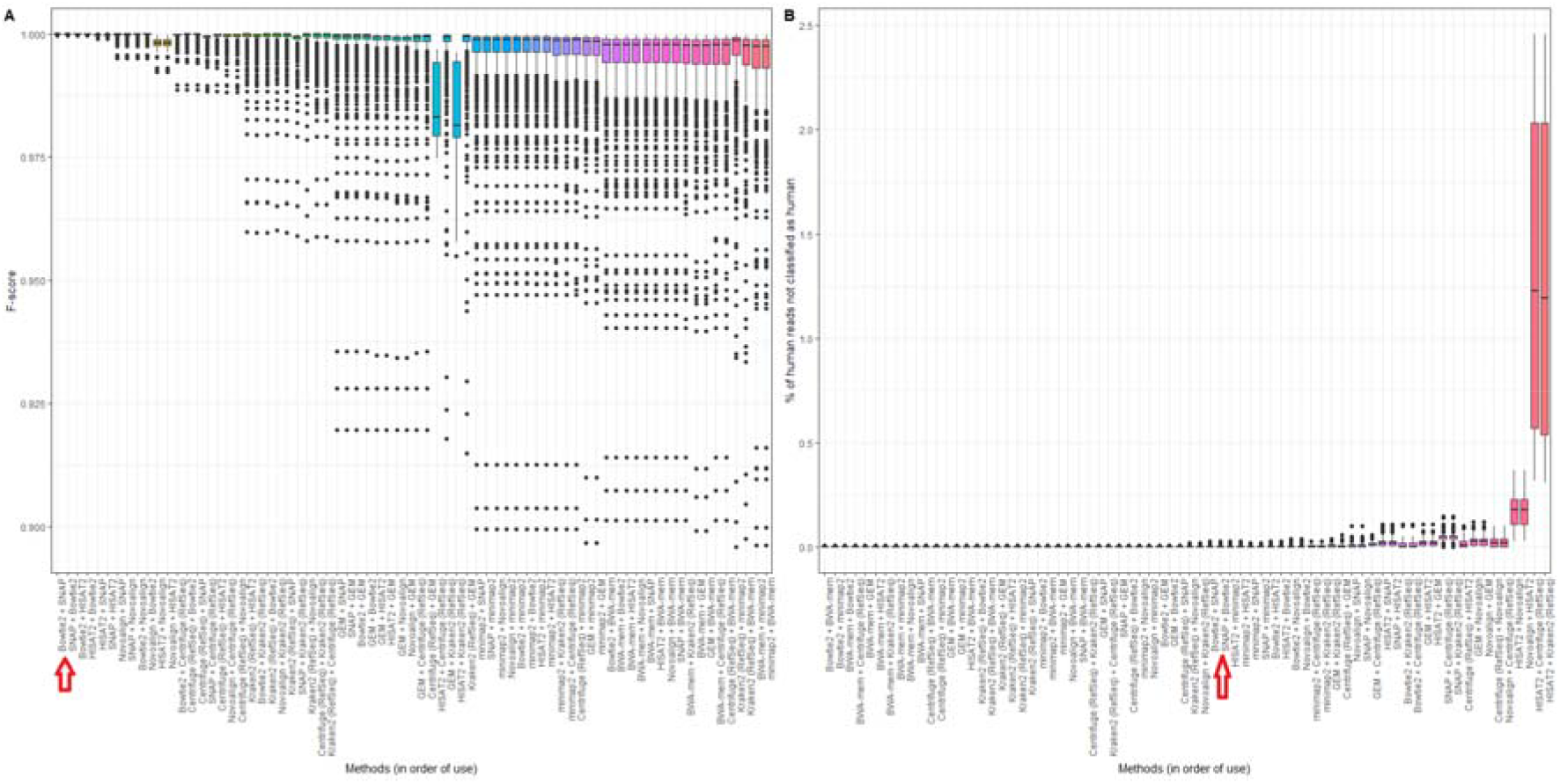
Performance of all 2-stage combinations of 9 independent methods of identifying human reads within a range of bacterial datasets (72 pairwise combinations; data comprises 10 bacterial species, each with between 1 and 10% simulated human contamination, using both 150bp and 300bp reads). All reads were simulated from, and where relevant aligned to, human genome version GRCh38.p12. The subfigures show (A) the F-score, and (B) the percentage of human reads not classified as human. Bars in both subfigures are ordered from left to right by increasing variance, and in alphabetical order for methods with equal variance. Bowtie2 + SNAP is indicated on the axis. Data for this figure is available in **Supplementary Table 7**.

While these results broadly recapitulate those of **Figure 1**, variance in the false negative rate was substantially reduced when combining methods, suggesting that different aligners could compensate for each other’s omissions. This was particularly apparent when combining the two individually poorest-performing aligners in terms of false negative rates in **Figure 1B**, Novoalign and HISAT, which reduced the median false negative rate of each aligner from approx. 1.5% to 0.18% (**Figure 5**) (although these were still by far the worst performing method for false negative rates, having little utility as the first method in a two-stage process).

The optimal combination of methods, in terms of F-score, was Bowtie2 followed by either SNAP or HISAT2. The F-score was 1 for 541 of the 600 Bowtie2/SNAP simulations and 416 of the 600 Bowtie2/HISAT2 simulations. Similarly, the false negative rate was 100% for 588 of the 600 Bowtie2/SNAP simulations and for 471 of the Bowtie2/HISAT2 simulations, although in both cases the absolute number of false negative calls, if any, was extremely low (< 10 reads; **Supplementary Table 7**).

### Does it matter if only most, but not all, human reads are removed?

In our simulations, relatively few human reads were retained in absolute terms. However, our simulations used relatively low sequencing depths of < 800,000 total reads for each of the 10 species (hence, < 80,000 true human reads in any one sample, as detailed in **Supplementary Table 1**). Although we found that all methods of human read depletion resulted in a consistently small proportion of retained human reads (typically < 0.1%), as sequencing depths increase (concomitant with decreasing cost) this could still be hundreds of human reads in absolute terms. Furthermore, these reads are unlikely to be randomly distributed throughout the human genome, as demonstrated above for the (repetitive) sex chromosomes. Other regions of the human genome may also have similar problems (although we did not directly observe this in our simulations). For instance, the human leukocyte antigen (HLA) on chromosome 6p21.3 is hypervariable, having > 10% sequence divergence between haplotypes (39). If using a ‘general purpose’ aligner, reads are highly unlikely to map to this region. As such, these reads would be disproportionately retained within a ‘human depleted’ bacterial dataset.

Assuming a sufficient number of reads across a given region, it is possible that variants within it could be called with confidence and used to impute phenotype (for instance, variants in HLA genes have well-documented associations with immune disorders (40)) and/or be associated with named individuals, assuming the identities of those involved in sample preparation were also known. Although anonymising samples is routine practice (and to some extent ameliorates this risk), generators of raw sequencing data frequently waive their own anonymity by virtue of publication. Furthermore, while human variant calls are often filtered on the basis of minimum depth (number of reads covering that position) and base call frequency (proportion of reads supporting a particular allele), in absolute terms this rarely represents large numbers of reads. For instance, the default recommendations applied by the variant calling pipeline NASP (41) are a minimum depth of 10 reads, of which 9 must support a given SNP.

To demonstrate that these are not just theoretical concerns, we reasoned that the highest-performing method, the Bowtie2/SNAP two-stage approach, would be able to best recover human reads inadvertently retained within public datasets. To test this possibility, we parsed the European Nucleotide Archive to obtain a diverse range of Illumina paired-end genome sequencing reads from each of the 10 bacterial species used in our simulations (see Materials and Methods). In total, we obtained 11,577 SRA sample IDs, representing sequencing data from 356 different BioProjects. From each sample we identified human reads using the two-stage method of Bowtie2/SNAP and then called SNPs – which could be used to predict individual phenotypic characteristics – using a BWA-mem/mpileup pipeline (see Materials and Methods). The results are summarised in **Table 2**, with the number of human reads detected per sample, and associated phenotypic predictions, given in **Supplementary Table 8** (in which sample and BioProject IDs are redacted).

While in absolute terms the number of human reads identified in the 11,577 samples was often low, there were several prominent outliers, with 58 samples (0.5%) containing > 10,000 human reads and 299 (2.6%) containing > 1000. Overall, 100 or fewer human reads could be detected in 8321 samples (71.8%), 10 or fewer human reads in 2550 (22.0%), and no human reads in 464 (4.0%). Across the 11,113 samples in which 1 or more human reads were detected, the mean number of reads detected was 1255. With each sample sequenced at a depth of 1-5 million reads, this represents 0.02-0.1% of the number originally sequenced, consistent with our expectation from simulations.

**Table 2.** SNPs called from contaminating human reads within publicly-archived sequencing datasets.

| Status of sample | No. of samples | % of samples |
| --- | --- | --- |
| Contains 1 or more SNPs | 7389 | 63.8 |
| Contains 1 or more SNPs with $\geq 2$ reads supporting alt allele, $QUAL \geq 20$, and $MAPQ = 60$ | 1657 | 14.3 |
| Contains 1 or more SNPs considered 'common' in dbSNP | 5113 | 44.1 |
| Contains 1 or more 'common' SNPs, each with $\geq 2$ reads supporting alt allele, $QUAL \geq 20$, and $MAPQ = 60$ | 731 | 6.3 |
| Contains 1 or more SNPs recorded in ClinVar | 412 | 3.6 |
| Contains 1 or more SNPs recorded in ClinVar, each with $\geq 2$ reads supporting alt allele, $QUAL \geq 20$, and $MAPQ = 60$ | 191 | 1.6 |
| Contains 1 or more 'common' SNPs recorded in ClinVar, each with $\geq 2$ reads supporting alt allele, $QUAL \geq 20$, and $MAPQ = 60$ | 13 | 0.11 |

It is possible to call SNPs even from a limited number of human reads (and so impute phenotype), although we do not expect the majority of calls to be made with high confidence. Without applying any filter criteria, one or more human SNPs could be called from the residual human reads in 7389 (63.8%) of samples (**Table 2**), although it is reasonable to believe a high error rate. We applied several post-processing criteria to retain only higher-confidence SNPs, requiring each alternative allele to be supported by 2 or more uniquely-mapped reads and to have been previously reported in a major human population (i.e. the SNP is considered ‘common’; see Materials and Methods). After applying these criteria, SNPs could still be called in 731 samples (6.3%) (**Table 2**).

From these higher-confidence SNPs, it was possible to impute particular phenotypes. For instance, in 13 samples (0.1%), one or more ‘common’ SNPs were called that were also recorded in the ClinVar database (42). While these SNPs were all considered ‘benign’ or ‘likely benign’ (**Supplementary Table 8**), this is likely because SNPs with pathological clinical significance are uncommon (i.e., this subset of ClinVar SNPs would not meet the dbSNP definition of ‘common’, which is based on minor allele frequency; see Materials and Methods). By relaxing this requirement, we found 191 samples (1.6%) with one or more SNPs supported by multiple uniquely-mapped reads and which were present in ClinVar but not also ‘common’ (**Table 2**). Within this subset, we found three samples with SNPs indicative of adverse drug responses (ClinVar variant IDs 12351, 17503 and 37340) and 19 samples with pathological SNPs, including hereditary predisposition to various cancers (including breast and ovarian), polycystic kidney disease and Leber’s optic atrophy (**Supplementary Table 8**).

These analyses of real data are broad in scope and intended to illustrate a general point: that phenotypes can be imputed from even a small number of reads. Overall, these results establish that phenotypically distinct human sequence is widespread in publicly archived bacterial data.

Nevertheless, for the majority of samples, a low number of SNP calls may only be minimally informative. As only a proportion of reads will contain SNPs, we can suggest a reasonably broad level of tolerance for the number of human reads retained within a real bacterial sample (although still advocating as exhaustive an approach to their removal as possible). Across all 11,577 samples, we made 12,215 higher-confidence SNP calls (those where the alternative allele was supported by 2 or more uniquely-mapped reads), of which 2063 were also ‘common’ (**Supplementary Table 8**), i.e. more likely to be annotated with phenotypic characteristics. We found that on a per-sample basis, fewer than 10 ‘common’ SNPs could be called with reasonable confidence from samples containing up to 10,000 human reads (**Figure 6**). While this suggests a relatively large number of human reads could be retained in a microbial dataset without being unduly informative, it is important to note that individual SNPs (which could still be clinically significant) could also be called with confidence even at far lower depths (< 100 reads; see **Figure 6**). Furthermore, there is already a substantial body of overt phenotypic associations with single SNPs, including, for instance, blue eyes (43), lactose intolerance (44) and alcohol-related flushing (45).

**Figure 6.**
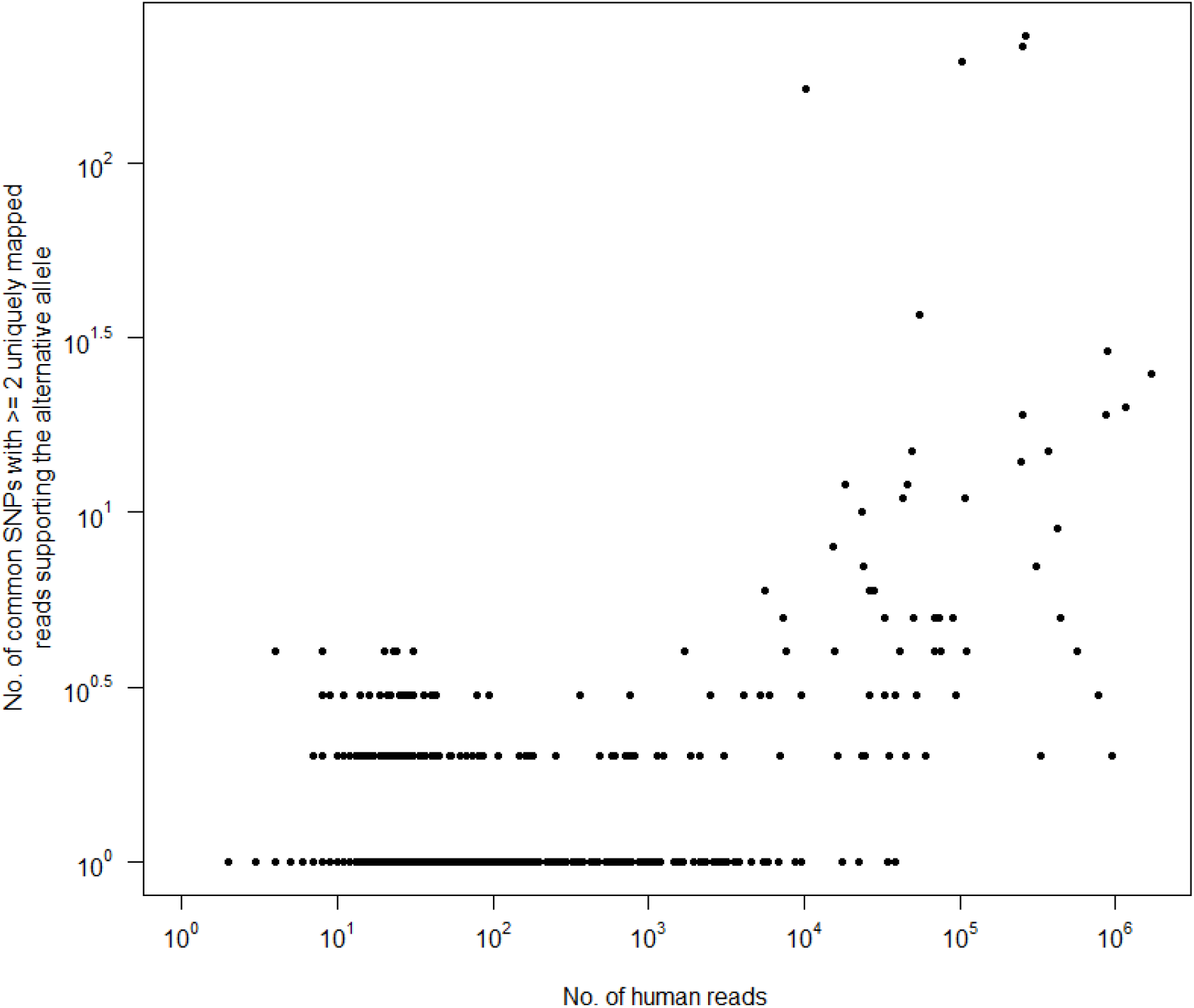
Relationship between the number of human reads retained within 11,577 publicly-archived bacterial readsets and the number of higher-confidence ‘common’ SNPs called using them (i.e. SNPs previously called by the 1000 Genomes Project in at least one of 26 major human populations, with the alternative allele supported by ≥ 2 uniquely mapped reads). Human reads were identified by aligning all reads to the human genome using Bowtie2 followed by SNAP. Data for this figure is available in **Supplementary Table 8**.

### Recommendations for depleting human reads from microbial datasets

If using shorter (150-330bp) bacterial reads, then to maximise the number of human reads whilst minimising the risk of false positive calls, we recommended using Bowtie2 (which performs near-exact read mapping, optimising parameters where appropriate to account for read length), followed by either of the aligners SNAP or HISAT2. Our analyses suggest that these aligners could complement each other and so the cost of using two methods sequentially would be incurred only in additional computational time, not in reduced accuracy. The individual methods with lowest false negative rates (that is, those which successfully detected the majority of human reads) were BWA-mem and Kraken2, although these methods both had relatively high false positive rates and so could erroneously classify bacterial reads as human. This was particularly apparent for genomes rich in mobile elements, such as *C. difficile*.

Classification accuracy was strongly affected by read length, particularly when using alignment-based methods (**Figure 3**). We found that when sequencing viral reads or bacterial reads of > 300bp, the most accurate, as well as simplest and quickest, approach was to use direct classification methods (Centrifuge and Kraken2). Although runtime was not formally assessed in this study, we noted that classification-based methods were far faster to complete than alignment-based human read detection. Importantly, and irrespective of the speed of a given aligner, ‘alignment’ is in practice a multi-step process, requiring the subsidiary steps of BAM sorting, BAM indexing, BAM subsetting and read ID extraction (that is, converting BAM to BED). This approach is intrinsically slower than the use of either Kraken2 or Centrifuge, and so these methods could be preferred for shorter reads if speed is a consideration: Kraken2 having the lower false negative rate, and Centrifuge the lower false positive rate (for longer reads, we found no discernible difference).

An additional benefit of Kraken2 and Centrifuge are that their databases can easily be customised to contain multiple human genomes rather than a singular reference (of which the current human reference is essentially an idiosyncratic type specimen (46)). This approach would expand the number of human k-mers (against which k-mers from the reads are compared) to include those from divergent regions, such as the HLA, and could also incorporate population-specific sequence (for example, recent deep sequencing of 910 individuals of African descent identified 296 Mb of sequence missing from the GRCh38 reference (47)). Reads originating from population-specific regions would not be identifiable using either of the approaches tested in this study. An additional complication with alignment-based, relative to classification-based, methods is that as divergence increases relative to a reference the number of reads correctly mapped to the reference will decrease, as previously demonstrated for influenza A (48). While in our simulations, viral reads (including from influenza A) could be consistently mapped to the same reference genome from which they were sequenced, we would not expect this to be as pronounced for non-simulated reads (that is, where the genome from which the reads were derived diverges from the reference).

Alignment-based methods can, however, allow a finer degree of discrimination than classification-based methods. An alternative alignment-based approach would be to align reads (that are nominally from one bacterium) not just to the human genome but to the human and bacterial references simultaneously, as in the ‘remove contamination’ module of Clockwork (http://github.com/iqbal-lab-org/clockwork), a pipeline developed for the CRyPTIC project to process reads from *M. tuberculosis*. Should reads have multiple mapping locations, this can allow finer discrimination between human and non-human sequence. If a read, or one mate of a pair of reads, maps both to the human and microbial genomes, the user can decide whether to consider those reads as human or microbial in origin, so maximising either the number of human (or human-like) reads removed or the number of microbial (or microbial-like) reads retained.

In terms of memory requirements, while both classifiers drew upon reasonably large databases (approx. 8Gb in each case), aligners also required a pre-built index of the (approx. 3Gb) human genome as input. These indices varied 10-fold in size, from 2.3 Gb (SMALT) to 4.0 Gb (Bowtie2), 4.3 Gb (HISAT2), 5.1 Gb (BWA), 6.9 Gb (Minimap2), 8.1 Gb (Novoalign), 12.9 Gb (GEM) and 28.9 Gb (SNAP). One of the highest-performing methods in this study, Bowtie2 followed by SNAP, accordingly had among the largest runtime and memory requirements.

Nevertheless, it is important to take an exhaustive approach to depleting human reads from bacterial datasets. This is because as variant databases increase in scope and volume, it should become increasingly likely that personally-identifiable phenotypic characteristics could be recovered from a small number of reads. This study has demonstrated that the subtractive alignment of human reads, if using only one aligner to do so, will likely be insufficient. Further to this, it remains a strong possibility that there is endemic, albeit low level, human read contamination of (older) bacterial datasets in public archives.

## Materials and Methods

### Simulating sets of human-contaminated microbial reads

We obtained the NCBI reference genomes (criteria for which is detailed at http://www.ncbi.nlm.nih.gov/refseq/about/prokaryotes/, accessed 16^th^ August 2018) for 10 clinically common species with fully sequenced (closed) core genomes, as detailed in **Supplementary Table 1**, alongside versions GRCh38.p12 and GRCh37.p13 of the human primary assembly (that is, the consensus genome excluding alternate haplotypes and patches; ftp://ftp.ensembl.org/pub/release-96/fasta/homo_sapiens/dna/Homo_sapiens.GRCh38.dna.primary_assembly.fa.gz and ftp://ftp.ensembl.org/pub/release-5/fasta/homo_sapiens/dna/Homo_sapiens.GRCh37.75.dna.primary_assembly.fa.gz, respectively; downloaded 13^th^ April 2019).

From each bacterial genome, 3 sets of 150bp and 300bp paired-end short reads (characteristic of the Illumina NextSeq and MiSeq sequencers, respectively) were simulated using dwgsim v0.1.11 (http://github.com/nh13/DWGSIM) with parameters-e 0.001-0.01 (non-uniform per-base error rate increasing across the read from 0.01 to 0.1%, approximating an Illumina error profile) and -y 0 (0% probability of simulating a random DNA read). All reads were simulated with an insert size of ((read length * 2) + (read length * 0.2)), i.e. 150bp reads were simulated with an insert size of 330bp. dwgsim does not output the otherwise randomly generated seed used for each simulation, although does allow seeds to be provided. To ensure results were reproducible, the same seeds were used for each of the three replicates of each read length: 123456789, 234567891, and 345678912, respectively. The number of reads sequenced for each genome is equivalent to a mean base level coverage of 10-fold (for bacteria) or 100-fold (for viruses), detailed in **Supplementary Table 2**.

To each of these microbial readsets was added a number of human reads equivalent to between 1 and 10% of the microbial reads, at increments of 1%. All human reads were simulated from either GRCh38.p12 or GRCh37.p13 and used the same dwgsim seeds and parameters as above. Notably, dwgsim assigns read names on the basis of their origin. In this way, we can easily determine which of a set of reads predicted to be either human or microbial are correctly classified: true microbial reads will have names corresponding to microbial chromosome IDs.

### Methods for detecting human read content

Two basic methods were used for detecting human reads within the contaminated microbial datasets: by alignment of all reads against the human genome, and by predicting read origin using the taxonomic classification tools Centrifuge v1.0.4 (19) and Kraken2 v2.0.7 (20). Alignments were performed using BWA-mem v0.7.17 (23), Bowtie2 v2.3.4.1 (49), GEM v3.6.1-16 (24), HISAT2 v2.1.0 (25), minimap2 v2.10-r773 (26), Novoalign v3.09.00 (www.novocraft.com), SMALT v0.7.6 (http://www.sanger.ac.uk/science/tools/smalt-0), and SNAP v1.0beta.18 (27), in all cases with default parameters. All alignments were made against 2 different builds of the human genome: the primary assembly of GRCh38.p12 (corresponding to GenBank assembly ID GCA_000001405.27 and obtained from Ensembl v96), and the primary assembly of GRCh37.p13 (corresponding to GenBank assembly ID GCA_000001405.14 and obtained from Ensembl v75).

In each case, BAM files were cleaned, sorted and duplicate-marked using Picard Tools v2.17.11 CleanSam, SortSam and MarkDuplicates, respectively (50). Supplementary and non-primary alignments were removed using SAMtools view v1.7 (51) with parameters -F 2048 -F 256. Finally, we obtained the set of reads mapped by each aligner (that is, considered human) using SAMtools view with parameters -F 4 -f 8, -F 8 -f 4, and -F 12, merging the resulting BAMs. This was in order to obtain reads where both ends mapped as well as the set of mapped reads with an unmapped mate, as the latter, being from the same DNA fragment, should also be of human origin. Read IDs were extracted from each BAM using the ‘bamToBed’ module of BEDtools v2.19.1 (52).

By contrast, the classification tools Centrifuge and Kraken compare sets of reads to a database of genomes, probabilistically assigning a taxonomic rank to each read. For both tools, we used one database containing only the human (GRCh38) genome and one containing the set of RefSeq bacterial, archaeal and viral genomes, plus the human genome. The human-only databases were custom-built using the same primary assembly from which human reads were simulated. For the larger multi-species databases, we obtained the pre-built Centrifuge ‘P+H+V’ database (ftp://ftp.ccb.jhu.edu/pub/infphilo/centrifuge/data/p+h+v.tar.gz, created June 2016) and the comparable MiniKraken (ftp://ftp.ccb.jhu.edu/pub/data/kraken2_dbs/minikraken2_v2_8GB_201904_UPDATE.tgz, created April 2019), both of which draw upon genomes stored in RefSeq.

### Human read detection in real sequencing data

To test for the presence of human reads in real bacterial sequencing data, we downloaded the daily-updated SRA BioProject summary file (n = 383,590 BioProjects; ftp://ftp.ncbi.nlm.nih.gov/bioproject/summary.txt, accessed 24th September 2019) and parsed it to extract a list of BioProject IDs with a data type of ‘genome sequencing’ and a species name matching, at least in part, the name of any of the 10 clinically common bacteria detailed in **Supplementary Table 1** (for example, ‘*Salmonella enterica* subsp. *enterica* serovar *Oranienburg* str. 0250’ matches ‘*Salmonella enterica*’). We then used the Entrez Direct suite of utilities (http://www.ncbi.nlm.nih.gov/books/NBK179288/, accessed 1st May 2019) to associate each BioProject ID with a list of SRA sample and run IDs (a ‘RunInfo’ file). RunInfo files were parsed to retain only those runs where ‘Platform’ was ‘ILLUMINA’, ‘Model’ (i.e. sequencer) was ‘HiSeq 2000’, ‘HiSeq 2500’ or ‘MiSeq’ (all of which use TruSeq3 reagent chemistry), ‘LibrarySource’ was ‘GENOMIC’ or ‘METAGENOMIC’, ‘LibraryStrategy’ was ‘WGS’, ‘LibraryLayout’ was ‘PAIRED’, ‘LibrarySelection’ was ‘RANDOM’, ‘avgLength’ was ≥ 150 (i.e., mean read length of 150bp) and ‘spots’ was > 1 and < 5 (i.e., approximating a read depth of > 1 and < 5 million reads, the upper limit chosen to minimise the computational cost of data processing). This generated a list of 11,577 SRA sample IDs, representing sequencing data from 356 different BioProjects.

From each sample, human reads were classified using the two-stage approach of Bowtie2 followed by SNAP (detailed above). These reads were extracted from the original fastq files using the ‘subseq’ module of seqtk v1.3 (http://github.com/lh3/seqtk) and aligned to the human genome (GRCh38.p12) using BWA-mem v0.7.17 with default parameters (BWA was chosen as it was among the most sensitive aligners tested). The resulting BAMs were post-processed using Picard Tools to clean and duplicate-mark (as above), with the subset of mapped reads obtained using SAMtools view v1.7 with parameter -F 4. SNPs were called using BCFtools mpileup (51) with parameters -A (do not discard anomalous read pairs, i.e. include singleton ‘orphan’ reads) and -B (disable probabilistic realignment for the computation of base alignment quality scores). These parameters were chosen so as to maximise the likelihood of SNP calling at extremely low depth, with the intention that lower-confidence calls could later be identified (see below).

The resulting VCF was annotated using BCFtools ‘annotate’ to assign dbSNP IDs, where available, to those positions already known to be *bona fide* human SNPs. For this purpose, we obtained two sets of known human SNPs: those with clinical assertions, which are included in ClinVar (ftp://ftp.ncbi.nlm.nih.gov/pub/clinvar/vcf_GRCh38/clinvar_20191013.vcf.gz, accessed 21^st^ October 2019), and the set of dbSNP ‘common SNPs’ (ftp://ftp.ncbi.nih.gov/snp/organisms/human_9606/VCF/common_all_20180418.vcf.gz, accessed 21^st^ October 2019). ‘Common’ SNPs are defined as having both a germline origin and a minor allele frequency of ≥ 0.01 in at least 1 of 26 major human populations (according to the 1000 Genomes Project (53); http://www.internationalgenome.org/category/population), with at least two unrelated individuals having the minor allele (http://www.ncbi.nlm.nih.gov/variation/docs/human_variation_vcf/, accessed 21^st^ October 2019).

Finally, higher-confidence human SNPs were considered those which had (a) ≥ 2 reads supporting the alternative allele (these counts were obtained from the DP4 field of the mpileup VCF, and automatically exclude low-quality bases [by default, those with Phred < 13]), (b) QUAL value ≥ 20, (c) MAPQ value of 60 (used by BWA to indicate unique mapping), and (d) could be assigned a ‘common’ dbSNP ID.

### Evaluation metrics

For each method of human read detection, we obtained the number of true positive (TP; human read classified as human), false positive (FP; bacterial read classified as human) and false negative (FN; human read not classified as human) read classifications. We then calculated the precision (positive predictive value) of each method as TP/(TP+FP), recall (sensitivity) as TP/(TP+FN), and as an ‘overall performance’ measure, the F-score (as in (30)), which equally weights precision and recall: F = 2 * ((precision * recall) / (precision + recall)). The F-score summarises the performance of each method as a value bound between 0 and 1 (perfect precision and recall).

### Gene ontology (GO) term enrichment

GO term enrichment was assessed using the R/Bioconductor package topGO v2.36.0, which implements the ‘weight’ algorithm to account for the nested structure of the GO tree (54). This is a closed testing procedure – the computation of p-values per GO term is conditional on the neighbouring terms, and so the tests are not independent. For this reason, p-values produced by the ‘weight’ algorithm are interpreted as corrected or not affected by multiple testing. topGO requires a reference set of GO terms as input. These were built from the GRCh38.p12 and GRCh37.p13 sets (obtained via Ensembl BioMart versions 96 and 75, respectively), each filtered to remove terms with evidence codes ‘NAS’ (non-traceable author statement) and ‘ND’ (no biological data available), and those assigned to fewer than 10 genes in total. Significantly enriched GO terms (p◻<◻0.05) were retained only if the observed number in each set of ‘false negative genes’ exceeded the expected number by 2-fold or greater.

## Supplementary Figures

**Supplementary Figure 1.**
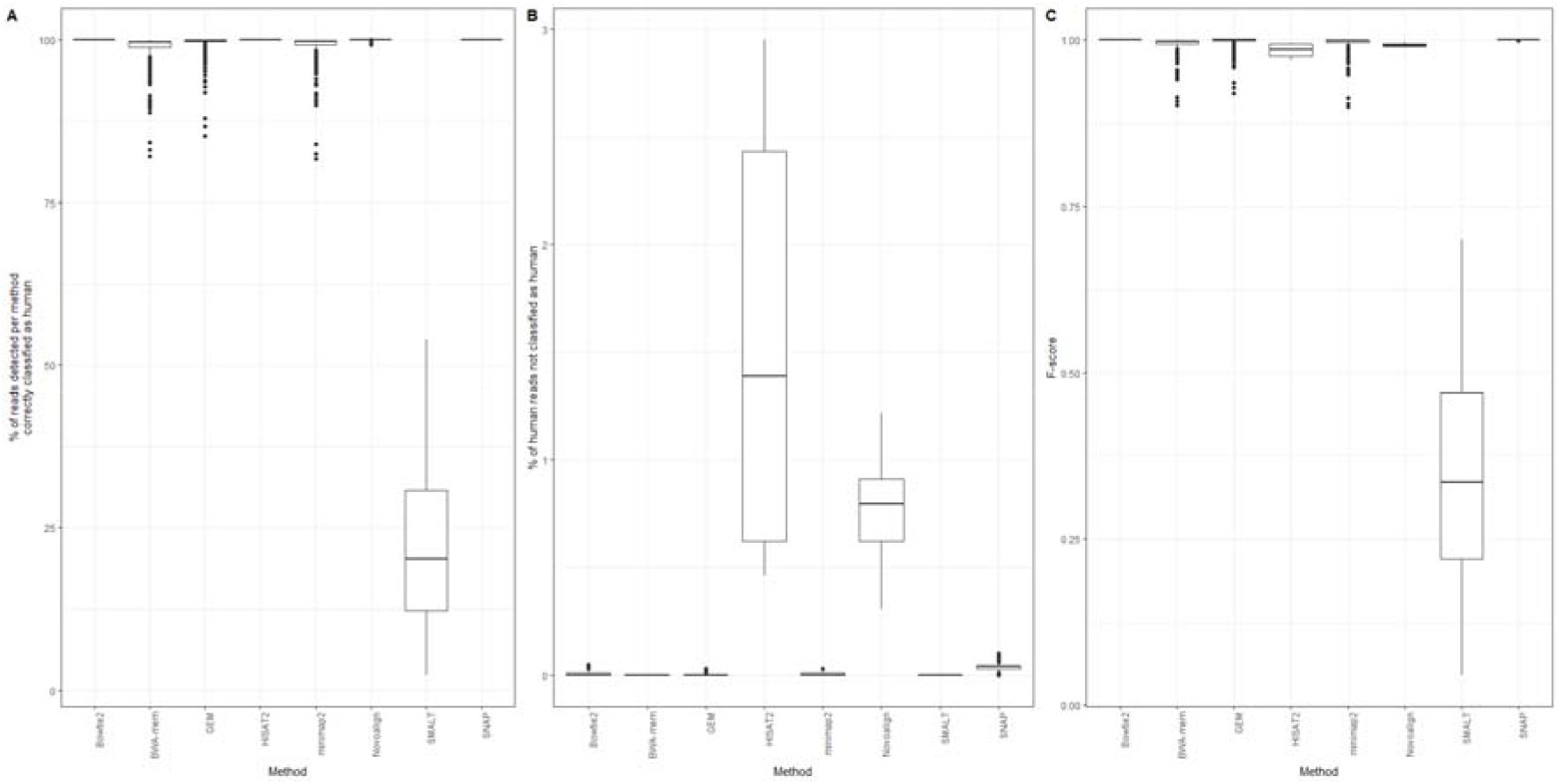
Performance of 8 different methods of identifying human reads within a range of microbial datasets (comprising 10 bacterial and 3 viral species, each with between 1 and 10% simulated human contamination, and sequenced using both 150bp and 300bp reads). All reads were simulated from human genome version GRCh38.p12 and aligned to a previous, lower-quality, assembly, GRCh37.p13. This was in order to approximate the effect of contaminating (human) sequences being in part degraded or incomplete and so not necessarily precisely mapping. The subfigures show (A) the percentage of reads correctly classified as human, (B) the percentage of human reads not classified as human, and (C) the F-score. Data for this figure is available in **Supplementary Table 2**.

## Supplementary Tables

**Supplementary Table 1.** Microbial genomes used in this study.

**Supplementary Table 2.** Performance of read classification and alignment methods at detecting human reads within microbial datasets.

**Supplementary Table 3.** Performance of read classification and alignment methods at detecting human reads within microbial datasets, when varying read length.

**Supplementary Table 4.** Number of human reads not classified as human by each method, and their location.

**Supplementary Table 5.** GO term enrichment among genes whose reads were not classified as human.

**Supplementary Table 6.** GO term enrichment among genes to which bacterial reads had been incorrectly assigned.

**Supplementary Table 7.** Performance of read classification and alignment methods at detecting human reads within bacterial datasets, after a 2-stage use of methods.

**Supplementary Table 8.** Phenotypically distinct human characteristics identified from reads contaminating bacterial datasets.

## Declarations

## Acknowledgements

The authors would like to thank Dr. Emily Clark (University of Edinburgh) for facilitating access to additional computational resources.

## Availability of data and materials

All data generated or analysed during this study are included in this published article (and its supplementary information files).

## Authors’ contributions

SJB conceived of and designed the study with support and supervision from TRC, TEAP, DWC and ASW. SJB performed all informatic analyses. SJB wrote the manuscript, with edits from all other authors. All authors read and approved the final manuscript.

## Consent for publication

Not applicable.

## Ethics approval and consent to participate

Not applicable.

