## Supplementary figures and images for "Phenotypically distinct human sequence is widespread in publicly archived microbial reads: an evaluation of methods for its detection"

### Supplementary Figure 1

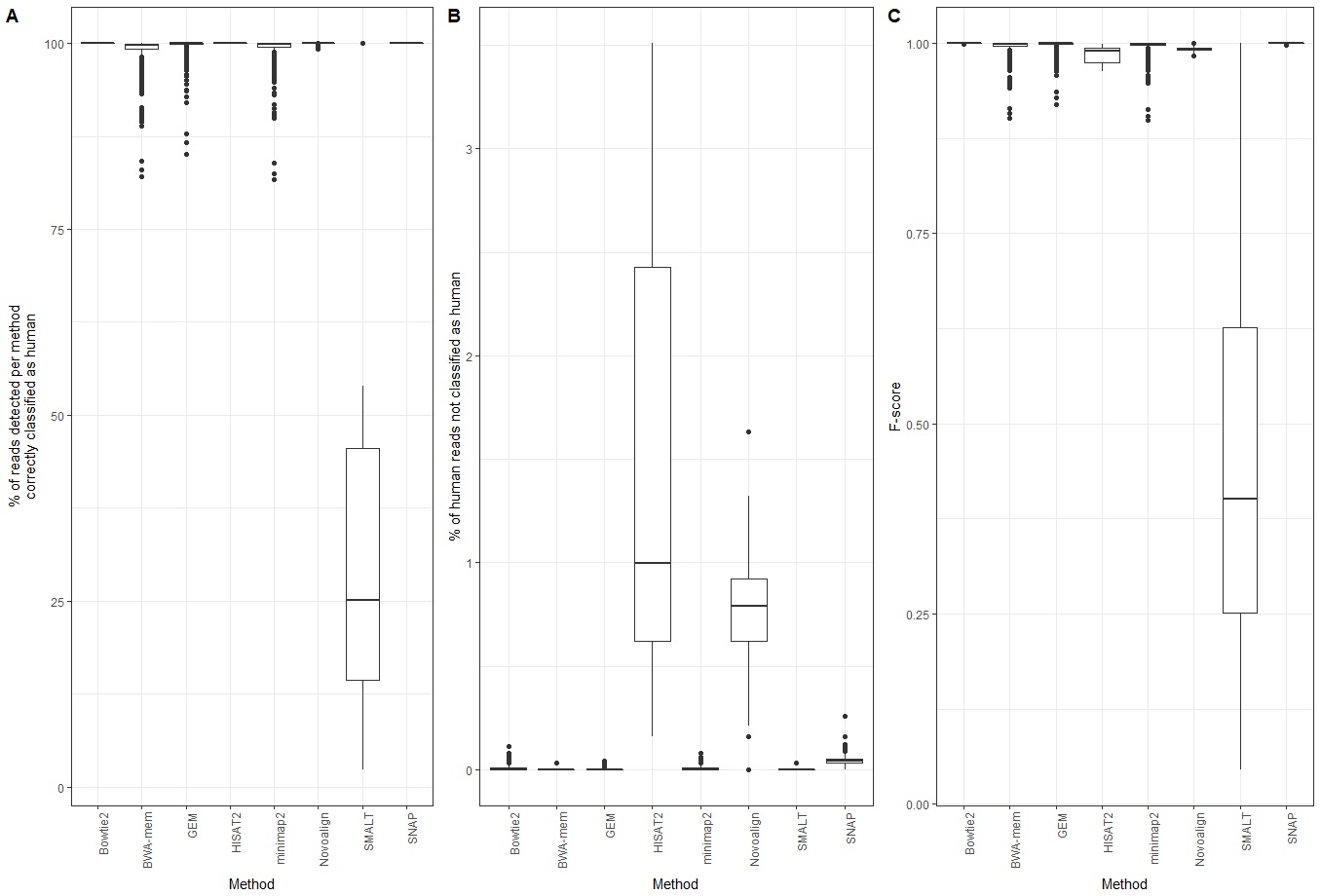
